# Ungulate personality and the human shield contribute to long-distance migration loss

**DOI:** 10.1101/2024.12.10.627781

**Authors:** Gavin G. Cotterill, Paul C. Cross, Eric K. Cole, Sarah R. Dewey, Benjamin L. Wise, Tabitha A. Graves

## Abstract

Long-distance ungulate migrations are declining and past research has focused on preserving migration paths where habitat fragmentation and loss disrupts movement corridors. However, changing residency-migration tradeoffs are the stronger driver of long-distance migration loss in some populations. The human shield effect relative to predation risk and anthropogenic food resources likely shapes these tradeoffs, but individual animals also vary in their propensity to tolerate proximity to humans and developed areas. We investigated how personality relative to human-habituation affects migration behavior. We categorized elk as bold or shy based on use of anthropogenic food resources identified through a clustering algorithm applied to GPS collar data. Bold elk were 4 times more likely to select wintering areas close to human activity and migrated 60% shorter distances compared to shy elk. As a result, elk wintering grounds were spatially structured such that conflict- and disease-prone individuals selected areas adjacent to human activity. Our results suggest that bold personality traits act as a precursor to human-habituation, which permits bold elk to reap the forage and predation rewards that occur in suburban landscapes. A multi-pronged approach beyond just maintaining habitat corridors may be necessary to conserve long-distance migrations for species that can become human-habituated.

## Introduction

Human development alters habitats and reshapes species communities, often by excluding species of higher trophic levels. Not all species are negatively impacted and, over time, those that have been displaced can return through natural recolonization or reintroduction efforts. Carnivores typically return last: they require both a sufficient prey-base and tolerance or policies that permit their presence (Cimatti et al., 2021). Thus, human-dominated landscapes create conditions which, through intentional policy decisions (e.g., hunting restrictions near homes) and unintentional circumstance (e.g., road density and predator road avoidance), limit predation risk from carnivores and humans alike. These processes (dubbed the human-shield effect; Berger 2007) can influence vital rates and create areas of high prey density and low predator density.

The same processes that give rise to the human shield alter incentives associated with movement. Without predators, sedentary movement by prey should reduce energetic costs if forage resources are adequate (Barker, 2018; Gower et al., 2008; Mitchell & Lima, 2002). In a suburban context, roads discourage movement (Passoni *et al*. 2021) and anthropogenic food resources create forage patches that are regularly replenished, which may disincentivize migration (Middleton et al., 2013; Mysterud, 2013) and increase pathogen transmission (Cross et al., 2007; Miller et al., 2003; Plaza & Lambertucci, 2017). Migration loss precludes benefits associated with the migration-escape hypothesis (Altizer et al., 2000), and carnivore exclusion limits the sanitizing effect of predators that selectively prey on infected animals (Brandell et al., 2022; Krumm et al., 2010; Packer et al., 2003). Human shielding thus creates multiple challenges: it can increase disease risk, contribute to migration loss, and generate direct human-wildlife conflict. Conflict is especially pronounced when individual animals become emboldened or food-habituated (Smith et al., 2023). Yet, habituation is a descriptive term encompassing innate and learned behaviors and cannot be easily identified in some contexts. As a result, management solutions can be opaque. Likewise, the processes by which individual animals come to exploit the human shield effect remain unclear.

Previous work demonstrated that suites of correlated behavioral traits among conspecifics are akin to personality (Sih et al., 2004), heritable (Réale et al., 2009), and can provide fitness advantages in specific contexts (E. F. Cole et al., 2012). For instance, fast-moving, bold individuals are more willing to engage with novel stimuli and, as a result, solve puzzles quicker than their slow-moving, shy counterparts (Benson-Amram & Holekamp, 2012; Mazza et al., 2018). But there may also be speed-accuracy tradeoffs such that shy individuals learn more slowly, but display greater flexibility when faced with changing conditions (Mazza et al., 2018; Sih & Del Giudice, 2012). Fitness mismatches also occur, as when incautious movements predispose individuals to capture (Wilson et al., 1993) or hunter harvest (Ciuti et al., 2012). Studies of captive and wild elk (*Cervus canadensis*) have placed individuals along a bold-shy personality gradient, wherein bold elk are more easily habituated to humans (Found, 2019) and bold individuals are more likely to abandon long-distance migration strategies (Found & St. Clair, 2019). Thus, individual elk likely vary across a behavioral spectrum in their ability to capitalize on the human shield effect. This, in combination with changing predator densities, climate, and development patterns may play a role in shaping human-elk conflict, migration loss, and disease dynamics.

The National Elk Refuge (NER), in Jackson, Wyoming, USA, is a 10,000 ha preserve that annually feeds ∼7,000 elk across 4 feeding areas. The placement of feeding areas subjects attending elk to a continuum of human presence: the northern boundary of NER abuts protected areas contiguous with Grand Teton (GRTE) and Yellowstone National Parks (YNP), while the southern boundary borders downtown Jackson, separated only by 8-foot fencing. Winter tourist activities are also confined to the southern two feeding areas. NER has fed elk in winter for over a century, and during that time, the Jackson herd has retained a mix of migratory tactics, even though currently >80% of the herd is fed for much of the winter. The recent arrival of chronic wasting disease to the Jackson elk herd prompted the U.S. Fish and Wildlife Service (FWS), which manages the refuge, to consider changes that include halting winter feeding operations and reducing elk numbers to mitigate the consequences of disease (Cook et al., 2024). One concern is that feed cessation could cause more elk to move into the suburbs during winter and thus spur conflict (Cotterill et al., 2024). In this, and other, mixed-migratory elk herds, higher recruitment among short-distance migrants was attributed to higher adult and calf survival (E. K. Cole et al., 2015) as the result of decreased predation risk (Hebblewhite & Merrill, 2011). Hunter harvest also disproportionately targets long-distance migrants that spend more time on public land. As a result, the proportional shift toward more resident and short-distance migration in the herd and additional elk-human conflict is likely to continue absent novel insights and harvest strategies. Our goal was to broadly evaluate the intersection of elk personality, conflict, and migration.

We identified concentrated patch use representing anthropogenic food resources using a clustering algorithm applied to global positioning system (GPS) collar data from 103 female elk. This type of cluster analysis has previously been used to identify ephemeral resources like kill-sites by predators (Clapp et al., 2021) or natal sites (Forshee et al., 2022). We classified individuals as bold if they generated clusters tied to anthropogenic food resources. We expected that individuals would generate clusters in proportion to the amount of time they spent in suburban areas and that boldness would be more frequent among elk migrating short distances. We also predicted that bold elk would spend more time on areas of the refuge with high human presence. If so, this would imply that elk personality plays a role in structuring elk space-use, selection of migration strategy, and would concentrate the most conflict- and disease-prone individuals on areas of the refuge with greater human presence.

## Materials and Methods

### Elk collar data

The dataset included 101 individual female elk monitored for at least a full winter period between 2016 and 2023. All GPS collars were programmed to record coordinates at 1.5-hour intervals, with most collars active for two to three years. All elk were collared on the NER during winter at three of four possible feeding areas using chemical immobilization or a baited corral trap, and most spent the bulk of their winters on the refuge.

### Cluster locations

We used the clustering algorithm from the package GPSeqClus (Clapp et al., 2021) in R (R Core Team, 2022) to identify foci representing extended, concentrated use in the suburbs. Elk contributed to the cluster analysis if they were monitored for at least 5 winter months, defined here as December 1 to April 30. The algorithm was applied to individuals’ winter movement trajectories. The minimum number of observations for a site to become a cluster was 8 locations, representing approximately 12 hours of use based on the 1.5-hour observation interval. We assigned points to a cluster if within 50 meters from a centroid during a time window of 3 days. The time window specified the maximum allowable period without visits before the cluster was considered abandoned. In other words, a cluster continued to accrue points so long as the individual returned to within 50 meters of its centroid every three days. Twelve hours of use in a 3-day period was deemed long enough to suggest some kind of reward, especially in developed areas where such use patterns were likely to be noticed. It may also have been conservative (shorter use periods would have created more clusters, some of them possibly associated with anthropogenic food), but yielded a feasible number to compare against known elk-depredation locations and to examine each cluster individually using satellite imagery.

We hypothesized that winter suburban clusters would disproportionately represent locations with provisions for livestock, including small hobby farms, horse properties and cattle operations. Managers receive and respond to landowner complaints concerning elk use of private property in winter, but landowners that enjoy elk presence are unrepresented in this dataset. From managers’ perspective, the latter also represents a source of potential or realized conflict because elk are large, mobile, and can cause damage while moving from or to these properties. Therefore, while not inclusive of all circumstances concerning to managers, we used the landowner complaints to assess the sensitivity of the clustering algorithm with respect to detecting anthropogenic food resource use in winter. We consulted managers from the Wyoming Game & Fish Department (WGFD) to determine if clusters corresponded to complaint locations. We also examined aerial imagery and documented whether clusters were within 100 meters of sheds, corrals, haystacks, or other confirmed complaint locations. We defined the suburbs as private lands surrounding Jackson, Wyoming, excluding part of the Fall Creek Herd Unit bounded by a major highway and unused by our collared elk.

### Migration classification

We classified individuals’ migration strategies following Cole *et al*. (2015) as short- or long-distance migrants. Short-distance migrants were those whose summer ranges occupied the private land complex and portions of Grand Teton National Park between Wilson and Moose, Wyoming. All other summer range categories were considered long-distance migrants. We also calculated the Euclidean distance between individuals’ mean March (winter) and July (summer) locations and evaluated both the categorical and continuous migration variables.

### Space-use while on the National Elk Refuge

We sought to identify potential spatial structuring of elk while attending the NER in winter. Anecdotally, managers noted that elk collared at the southern-most feeding area (adjacent to downtown Jackson and where more human activity occurs) tended to display unique behaviors like using different winter ranges from one year to the next. Our hypotheses included that elk might select feeding areas (1) at random; (2) on a first-come-first-serve basis where elk arriving earlier in the season would select feeding areas with less human activity; (3) based on their migration strategy or distance; or (4) according to the degree to which they exhibited bold behavior, which we assumed would be highest among suburban food-subsidized elk or elk that used the suburbs more in winter.

Although feeding areas on the refuge are kilometers apart from one another, most elk visited multiple feeding areas during a winter. Additionally, an 8-foot fence on the NER perimeter forces elk to enter and exit the refuge from the north. Thus, to reach the southern-most feeding area, they must pass several others. To address this, we used Dirichlet regression to model proportional feeding area use as a function of boldness, arrival times, and migration strategy, with a random intercept of individual for those monitored >1 winter (Cotterill and Graves 2024). We defined boldness as a categorical variable where individuals that were ever observed to generate a winter cluster in the suburbs were labeled as bold and if not, shy. We separately considered the proportion of each winter an elk spent in the suburbs. Arrival time was the day of the biological year on which they first arrived at the refuge (where May 1^st^ = 1). We tested categorical migration classifiers (resident, short distance, etc.) as well as migration distance, which we calculated as the average Fall and Spring migration distance of a corresponding winter when both were available. We used the ‘brms’ package in R (Bürkner, 2017). Where *y_i_* is a vector of proportion values that is an observation of a random variable Y and *FA* is the feeding area,

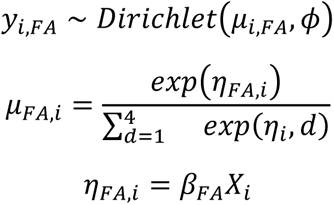

The link between *µ* and *η* is the multinomial function. *β_FA_* is a vector of slopes multiplying the predictor matrix *X*. *µ_FA_* priors were Student-T distributed (3, 0, 2.5). *ϕ* took a Gamma prior (0.01, 0.01). We fit the univariate models and an additive model including the three strongest predictors from the univariate model results. We assessed model predictive accuracy using leave-one-out cross validation (Hastie et al., 2009) and calculated the expected log-predictive densities (ELPD) as a relative index of predictive performance (Gelman et al., 2014). We report the differences in models’ ELPD and the standard error of ELPD differences (Sivula et al., 2022). We ran all models with 4 chains and 1000 iterations after warm-up. We visually inspected trace plots, ensured all Ȓ <=1.1 (Gelman & Rubin, 1992) and examined effective sample sizes to assess convergence.

## Results

### Migration classification and clusters

We classified 101 elk as 28 short-distance (mean = 12.4 km, sd = 2.9 km) and 73 long-distance migrants (47.8, 19.3; Cotterill and Graves 2024). Of the total sample, 59 elk used the suburbs at some point during winter and 32 individuals generated 359 winter clusters in the suburbs. Of these, 167 clusters (47% of total) corresponded to anthropogenic food sources in the region as confirmed by landowner complaints made to the Wyoming Game and Fish Department. Additionally, 68 of the unconfirmed clusters (19%) were within 100 meters of livestock sheds or corrals visible from satellite imagery. All known complaint locations were identified by the clusters. Many of the unknown locations appeared to be associated with residential landscaping, golf courses, or may simply have been bedding locations.

### Space use while on the National Elk Refuge

Boldness, which we defined as whether elk generated winter clusters in the suburbs, was the strongest predictor of refuge feeding area selection (Table 1). The boldness-only model had the equivalent support to the more complex model including boldness, migration classification, and proportional winter use of suburbs. ELPD differences less than 4 are typically considered small, indicating that the models including boldness had similar predictive power. However, the more complex model made nearly identical predictions because the effect sizes of the other two variables were small. We present predictions from the parsimonious boldness-only model. All models converged with Ȓ = 1.

**Table 1.** Boldness was the strongest predictor of feeding area selection while on the National Elk Refuge as measured by differences in the expected log-predictive densities (ELPD_diff_) and the standard error of ELPD differences (SE_diff_) between Dirichlet models.

| Model | $ELPD_{diff}^*$ | $SE_{diff}$ |
| --- | --- | --- |
| y ~ Bold. + Sub. Use + Mig. Clas. | 0.0 | 0.0 |
| y ~ Boldness | -0.3 | 3.8 |
| y ~ Suburban Use | -13.4 | 7.1 |
| y ~ Migration Classification | -19.3 | 8.2 |
| y ~ Migration Distance | -23.3 | 10.5 |
| y ~ Null | -37.3 | 12.1 |
| y ~ Arrival Time | -40.7 | 11.9 |

\* $ELPD$ differences less than 4 are considered small, indicating that the two models including boldness as a variable had similar predictive power.
Abbreviations: $ELPD_{diff}$ , difference of expected log-predictive densities; $SE_{diff}$ , the standard error of $ELPD$ differences; Bold., Boldness; Sub. Use, Suburban use; Mig. Clas., migration classification.

Predictions from the boldness-only model estimated that bold elk spend 38% of their time on feeding areas at Headquarters (95% CI 32%-44%; Figure 1a), 43% at Nowlin (36%-49%), 15% at Poverty Flats (11%-21%), and 4% at McBride (3%-5%). By contrast, shy elk were predicted to allocate 11% of their feeding time to Headquarters (9%-13%), 35% to Nowlin (30%-40%), 48% to Poverty Flats (42%-54%), and 6% to McBride (5%-8%). None of our study elk were captured at McBride, thus its use is likely underrepresented in both groups. However, shy elk in our sample still selected it more frequently than bold elk.

**Figure 1.**
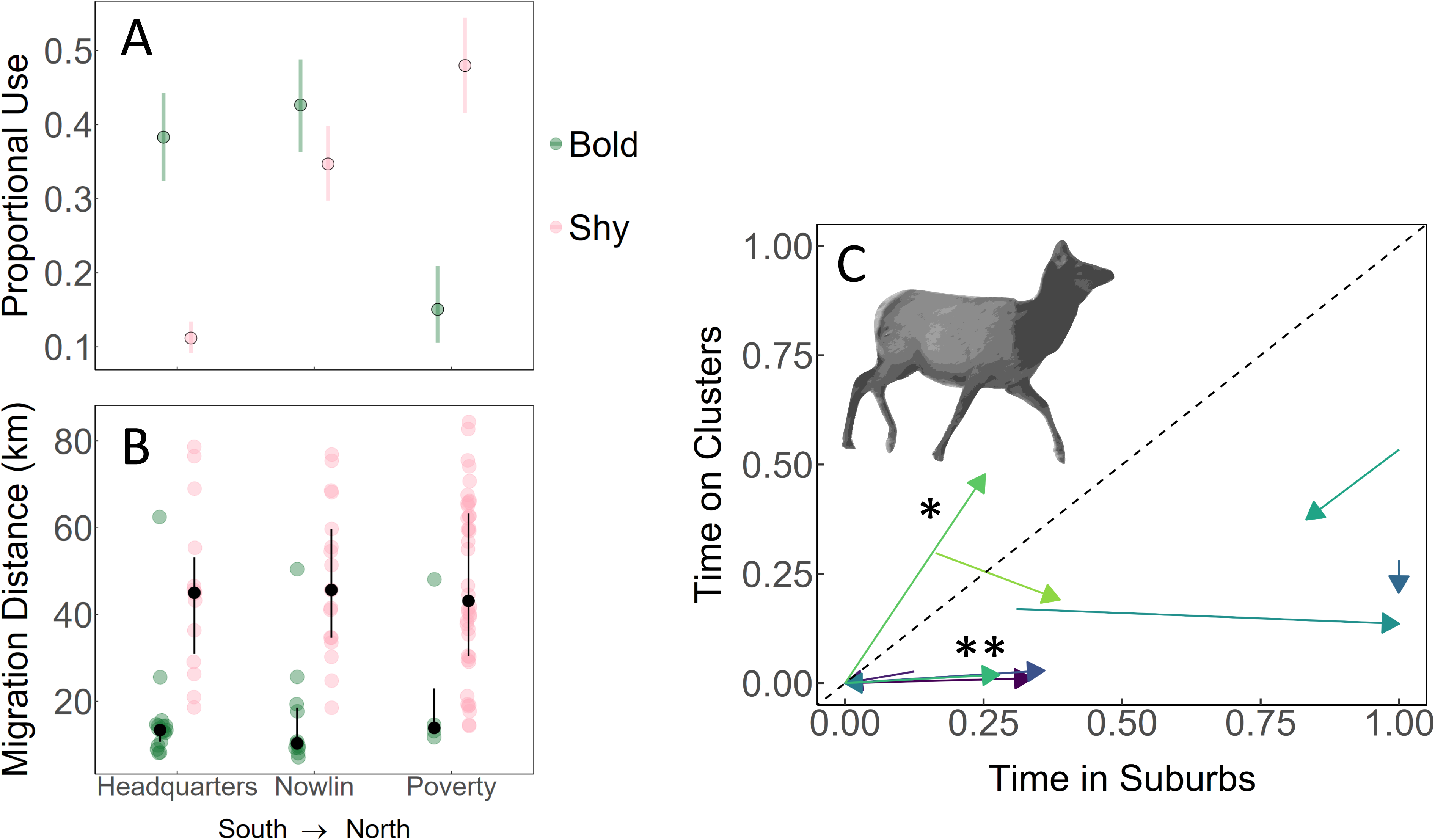
(A) Predicted proportional feeding area use of bold (green) and shy female elk (pink). 95% equal tail probability intervals denoted by error bars. (B) Spring migration distance (y-axis), and preferred feeding area while on the National Elk Refuge (x-axis) for bold and shy elk. Black points and lines represent the median and 25th and 75th percentiles. (C) Interannual changes in elk behavior relative to anthropogenic food use. Each arrow represents an individual monitored in two consecutive winters. Arrow origins represent the first winter; arrow terminuses, the following winter. The dashed black line represents a 1:1 relationship wherein time spent on clusters (anthropogenic food; y-axis) was a function of time spent in the suburbs (x-axis). Some elk, while in the suburbs, spent nearly half that time on clusters (*). Other elk spent significant time in the suburbs but rarely or never formed clusters (**). (art credit: Gavin Cotterill)

Bold elk migrated less than half as far (on average <20 km; Figure 1b) as shy elk (on average >40 km). However, migration strategy and distance were poor predictors of winter feeding area selection. Additionally, individual elk varied in their use of anthropogenic food in the suburbs, and of the suburbs more broadly, from year to year (Figure 1c). Most bold elk (but not all) tended to migrate shorter distances and select southern feeding areas with greater human presence (Figure 2a). Shy elk were more likely to be long-distance migrants, less likely to use the southern-most feeding area and less concentrated overall (Figure 2b). However, each feeding area hosted a mix of migration strategies. As a result, migration strategy was a poor predictor of refuge space-use (Table 1).

**Figure 2.**
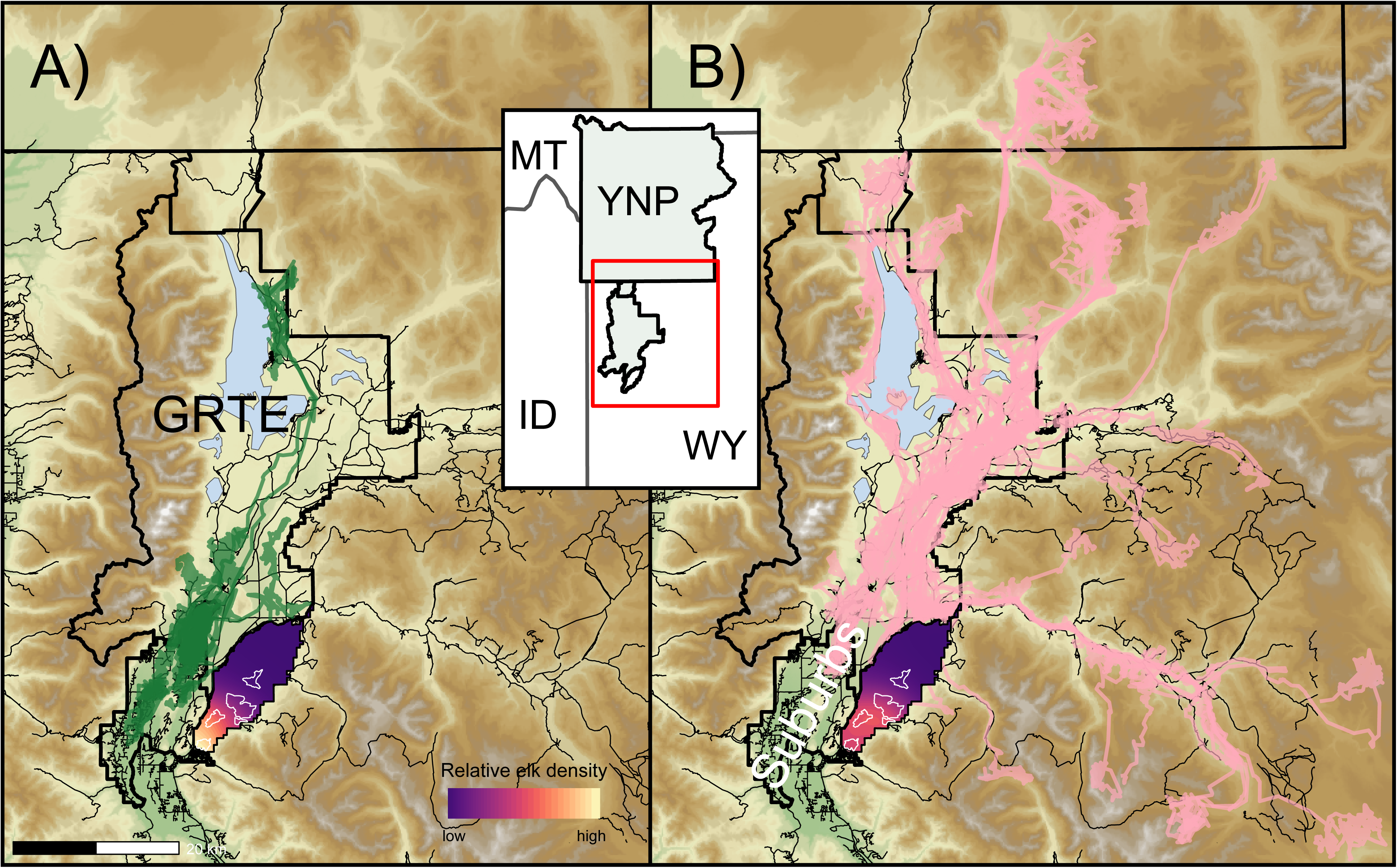
Elk that clustered on anthropogenic food sources in the suburbs during winter were classified as bold, otherwise shy. 75 randomly selected spring migration routes of bold (green paths; panel A) and shy (pink paths; panel B) elk and density plots illustrating relative space-use while on the National Elk Refuge (NER). Bold elk disproportionately used southern areas of the NER in closer proximity to human activity. Feeding areas outlined in white. Compared to bold elk, shy elk used areas north and east, farther from human activity and were less concentrated overall.

## Discussion

Our findings shed light on the capacity for wildlife to adapt to human-dominated landscapes and suggests a link between elk personality, behaviors relevant to forage-predation tradeoffs, and the persistence of long-distance migration. Bold elk, which we defined as those that formed suburban winter clusters, were 4 times more likely than shy elk to select the headquarters feeding area (adjacent to downtown Jackson) and migrated less than half as far. The combination of anthropogenic food sources and human-shield effect may work against managers trying to preserve long-distance migration. As long as the suburbs provide a safe haven to elk, the subset of the population able to exploit its lower predator densities and anthropogenic food subsidies will likely expand and constitute a larger share of the overall population.

While the suburbs may shield elk from predators, the same is not true for pathogens, particularly with respect to chronic wasting disease (CWD). Suburban spaces can support high Cervid densities that facilitate both direct and indirect CWD transmission while precluding the potential sanitizing effect of predators (which disproportionately prey on CWD-infected animals; (Brandell et al., 2022; Krumm et al., 2010). Compared to long-distance migrants, short-distance migrant elk spent more time concentrated in small areas, both on and off the refuge, that may become heavily contaminated with prions. CWD arrival in the Jackson Elk Herd could alter the fitness tradeoffs associated with these migration tactics and disproportionately harm bold elk if CWD infection risk is greater in suburban areas.

By applying the clustering algorithm to elk GPS data, we identified landowner complaint locations with 100% sensitivity. Bold elk disproportionately selected feeding areas closer to human activity. Short-distance migrants were more likely to generate suburban winter clusters than other migration strategies. However, not all short-distance migrants created these clusters, and some elk of both migration strategies did. Boldness was not merely a recapitulation of migration strategy or whether an elk used suburban areas at all, as evidenced by the lack of predictive accuracy in the migration classification models. Likewise, the fact that boldness outranked the proportion of winter an elk spent in the suburbs indicates that the manner of use (i.e., using anthropogenic food resources), rather than absolute use, was the stronger predictor of feeding area selection. Individual behavior also varied from year to year, with clusters being more likely in severe winters. Together, this suggests that during severe winters bold elk are more likely to seek out an easy meal in the suburbs. Our findings are consistent with behavioral research wherein bold elk were more prone to becoming human-habituated (Found & St. Clair, 2016). That same research concluded that bold elk are more likely to switch to resident migratory tactics; however, our elk were not monitored long enough to document trends toward residency over years and to-date there are no documented occurrences of this in the Jackson elk herd.

In the broader context of personality research, boldness in elk may represent individuals’ capacity to correctly identify the specific contexts in which reduced fear of humans yields advantage. This skill may result from a combination of innate and learned behaviors (Tablado & Jenni, 2017). If so, then bold and shy elk should occur across the landscape and different migration strategies. However, given stable conditions and specific rewards associated with the human shield, bold elk may come to make up a larger proportion of the population as recruitment among bold elk outpaces that of shy elk. The ideal dataset for testing these hypotheses would include cradle-to-grave tracking, known lineages, and measure physiological responses to certain stimuli. Nevertheless, we argue that simpler metrics based on movement patterns from GPS tracking may be adequate for some applications (Box 1).

Most elk collared in our study were darted along feedlines. Previous work showed that bold elk were more likely to be on the periphery of a herd and had shorter flight-initiation distances when approached (Found, 2019). Bold elk may therefore be more accessible to dart during capture. If bold elk are more likely to migrate short distances, this capture protocol may underestimate the proportion of long-distance migrants in a herd. In our dataset, darted elk were twice as likely to be classified as bold compared to those captured in a corral trap (*Χ^2^ p* = 0.07). However, the corral trap was only used on the feeding area most strongly associated with shy elk. With this confounding effect in mind, we make no inference with respect to randomness of our sample (or lack thereof) but take this as possible evidence of the role of personality in anthropogenic subsidy use and suggest that future work carefully consider the method of capture during study design.

Long-distance migrants in this system spend more time than their suburban counterparts on public land during hunting season. Hunting therefore disproportionately targets elk that represent a conservation priority. By contrast, suburban elk are often referred to as being outside managers’ reach, as hunting options there are limited. The patterns of space-use we identified of conflict-prone elk may allow managers to target this subset of the elk population while on the refuge, albeit imprecisely. Suburban space-use by Jackson elk is a relatively new phenomenon that has increased significantly in recent decades. Mechanisms driving this emergent behavior likely mirror those in other urban systems. If so, then the mix of increased social tolerance, reduced hunting and predation risk, and selection for bolder individuals may be creating conditions favoring higher recruitment among suburban residents (Breck et al., 2019; Hebblewhite & Merrill, 2011).

Most efforts to preserve long-distance migration routes have centered on conserving specific tracts of land. This is undoubtedly important work, as roads, fences, and human development can create significant barriers to wildlife movement. However, these efforts alone may fail without also limiting anthropogenic food sources and the strength of the human-shield effect. The methods employed here could help identify links between behavior and space-use that provide additional opportunities to support long-distance migration through alternative management approaches.

## Acknowledgments

Thanks to Kyle Lash (Wyoming Game and Fish Department) for his assistance with cluster validation. Funding for elk GPS collars was provided by the Grand Teton Association. Elk captures and handling occurred in compliance with Wyoming Game and Fish Chapter 33 permit number 394. Any use of trade, firm, or product names is for descriptive purposes only and does not imply endorsement by the U.S. government. The findings and conclusions in this article are those of the author(s) and do not necessarily represent the views of the U.S. Fish and Wildlife Service. This article has been peer reviewed and approved for publication consistent with USGS Fundamental Science Practices (https://pubs.usgs.gov/circ/1367/). The U.S. government is authorized to reproduce and distribute reprints for government purposes notwithstanding any copyright notation herein. The U.S. government retains, and the publisher, by accepting the article for publication, acknowledges that the U.S. government retains a nonexclusive, paid-up, irrevocable, worldwide license to publish or reproduce the published form of this paper or allow others to do so, for U.S. government purposes.

## Author contributions

GC and TG identified the modeling framework with substantial input from co-authors and wrote the initial draft; GC performed analyses. All authors contributed to revisions and gave final approval for publication.

## Data Availability Statement

Elk GPS data are available upon request via the National Elk Refuge and Grand Teton National Park

## Box 1: Key points relevant to conflict and migration management.

- Human-habituated animals are predisposed to conflict.
- Previous work demonstrated that individual elk vary in their capacity to become human-habituated and could be placed along a personality spectrum according to this and other shared traits.
- We labeled elk exhibiting behavior consistent with the use of urban food sources as “bold,” and otherwise, “shy.”
- Bold elk were more likely to migrate short distances and selected winter refuge areas with more human activity.
- Bold elk may be more easily captured by chemical immobilization (darting) than shy elk.
- Bold individuals may have less fear of humans in some contexts and therefore a greater likelihood of conflict than shy individuals.
- The degree to which certain behaviors are innate or learned remains unclear. Nevertheless, recognizing these patterns may help managers develop innovative ways to curtail conflict and preserve long-distance migration.

